# MOSim: bulk and single-cell multi-layer regulatory network simulator

**DOI:** 10.1101/421834

**Authors:** Carolina Monzó, Maider Aguerralde-Martin, Carlos Martínez-Mira, Ángeles Arzalluz-Luque, Ana Conesa, Sonia Tarazona

**Affiliations:** Genomics of Gene Expression Lab, Institute for Integrative Systems Biology, Spanish National Research Council (CSIC-UV), Paterna, 46980, Spain; Applied Statistics, Operational Research and Quality Department, Universitat Politècnica de València, València, 46022, Spain; Biobam Bioinformatics S.L., València, 46024, Spain

**Keywords:** Multi-omic simulator, bulk, single-cell, transcriptomics

## Abstract

As multi-omics sequencing technologies advance, the need for simulation tools capable of generating realistic and diverse (bulk and single-cell) multi-omics datasets for method testing and benchmarking becomes increasingly important. We present MOSim, an R package that simulates both bulk (via mosim function) and single-cell (via sc_mosim function) multi-omics data. The mosim function generates bulk transcriptomics data (RNA-seq) and additional regulatory omics layers (ATAC-seq, miRNA-seq, ChIP-seq, Methyl-seq and Transcription Factors), while sc_mosim simulates single-cell transcriptomics data (scRNA-seq) with scATAC-seq and Transcription Factors as regulatory layers. The tool supports various experimental designs, including simulation of gene co-expression patterns, biological replicates, and differential expression between conditions.

MOSim enables users to generate quantification matrices for each simulated omics data type, capturing the heterogeneity and complexity of bulk and single-cell multi-omics datasets. Furthermore, MOSim provides differentially abundant features within each omics layer and elucidates the active regulatory relationships between regulatory omics and gene expression data at both bulk and single-cell levels.

By leveraging MOSim, researchers will be able to generate realistic and customizable bulk and single-cell multi-omics datasets to benchmark and validate analytical methods specifically designed for the integrative analysis of diverse regulatory omics data.

**Key Points:**

1. MOSim is capable of generating synthetic datasets for a broad spectrum of omics types, supporting bulk RNA-seq, ChIP-seq, ATAC-seq, miRNA-seq, Methyl-seq, and transcription factor data, as well as single-cell omics, including scRNA-seq, scATAC-seq, and transcription factors.
2. MOSim enables the robust simulation of complex, many-to-many regulatory relationships across molecular layers, faithfully capturing intricate regulatory patterns.
3. Offering extensive options for customization, MOSim’s flexible experimental design and parameterization empowers users to simulate count matrices and multilayer regulatory networks, tailoring simulations to diverse experimental scenarios and omics types.

## Introduction

Rapid advancements in massive sequencing technologies have significantly facilitated the widespread adoption of multi-omic assays, enabling a comprehensive exploration of the regulatory mechanisms governing biological systems. Consequently, numerous bioinformatics tools have emerged to assist researchers in processing multi-omics data, with a specific focus on unravelling multi-layer gene regulatory networks (GRNs) [1,2]. These GRNs serve as interpretable computational models, providing insights into the intricate regulation of gene expression through interconnected networks. Notably, GRNs encompass diverse regulatory components, including transcription factors (TF), chromatin accessibility, long non-coding RNAs, micro-RNAs, and methylation, among others [2]. Despite the experimental capacity to generate both bulk and single-cell multi-omic sequencing datasets, a significant challenge in GRN studies lies in precisely integrating these multiple omic layers. Therefore, the importance of benchmarking, tuning, and validating multi-omics integration pipelines becomes evident.

Synthetic data, serving as ground truth, provides an indispensable resource for defining true positive and negative features sets, enabling rigorous benchmarking, tuning, and validation of analytical methods. Despite the paramount role of synthetic data, there are few publicly available algorithms capable of simulating multiple omic data types. To our knowledge, only three methods support comprehensive multi-omics simulation of gene expression regulation for bulk datasets. The first, the InterSIM R package [3], generates datasets for DNA methylation, gene expression, protein abundance, and their relationships. Although the method allows for customization of the number of biological replicates and the proportion of differentially expressed features, it lacks options for time series simulation and fails to report the interaction among features. The second tool, OmicsSIMLA C++ [4], can simulate genomics, transcriptomics, methylation, and proteomics data. Nevertheless, it restricts the generation of count data matrices to the transcriptomics module and does not include customizable options for time points or replicates. The third tool, the sismonr R package [5], simulates RNA-seq count data in conjunction with pre- and post-transcriptional regulatory networks, offering time-series simulation capabilities. Nonetheless, this method lacks the flexibility to customise expression profiles and dynamics, and the only omic quantification data it generates is gene expression.

Given the cell-type-specific nature of regulatory regions, it is surprising that only two methods currently support multi-omics simulation for single-cell datasets. The statistical simulator scDesign3 [6] encompasses scRNA-seq, scATAC-seq, CITE-seq and methylation. Meanwhile, scMultiSim [7] can simulate scRNA-seq and scATAC-seq datasets. While both methods accurately simulate datasets closely resembling real data, none of them provide essential customization options, such as the number of experimental groups, biological replicates, differentially expressed genes, accessible chromatin, and reporting of interaction between features. Importantly, none of these tools is designed to simulate gene regulatory relationships across omics features, which underscores the existing gaps and limitations in current multiomics simulation tools. GRouNdGAN [8] partially addresses this limitation by modeling GRNs with genes and TFs with single-cell resolution. However, it does not support other omic modalities, multiple experimental conditions, or multiple samples, further underscoring the need for more comprehensive simulation tools.

Here we present MOSim, a multi-layer regulatory network simulator for both bulk (RNA-seq, ATAC-seq, miRNA-seq, ChIP-seq and Methyl-seq) and single-cell datasets (scRNA-seq and scATAC-seq), implemented as an R Bioconductor package. In a nutshell, MOSim generates quantification data for each omics layer, precisely controlling active regulatory relationships between regulatory omics and gene expression data for differentially expressed genes. Moreover, MOSim empowers users to customise data generation, enabling the inclusion of experimental groups, biological replicates, time series, and diverse cell types. By harnessing the capabilities of MOSim, bioinformatic tool developers will be able to generate realistic and customizable bulk and single-cell multi-omics datasets, facilitating the benchmarking and validation of analytical methods tailored explicitly for integrating multi-omics data and inference of multi-layer GRNs.

## Results

### Overview of MOSim’s workflows

MOSim is a bulk and single-cell simulation environment designed for generating multi-omic regulatory networks with precise control over regulator-gene relationships. To create a synthetic ground truth multi-omic dataset, MOSim requires as input the list of omic data types to be simulated, a single sample of seed count data for each of them, and an association file for each regulatory omic type, indicating the *a priori* or potential regulatory features associated with each gene (Figure 1A). While MOSim provides users with example multi-omics datasets to use as seed count data for simulation, the algorithm may also be fed with the user’s count dataset of choice, regardless of organism, disease or platform of origin. Besides simulation of RNA-seq or scRNA-seq data depending on the type of study (i.e. bulk or single-cell), currently supported omic regulatory data types include ChIP-seq, miRNA-seq, Methyl-seq, ATAC-seq and scATAC-seq. The algorithm also supports modelling Transcription Factor (TF) - target gene interactions from both bulk and single-cell RNA-seq data.

**Figure 1:**
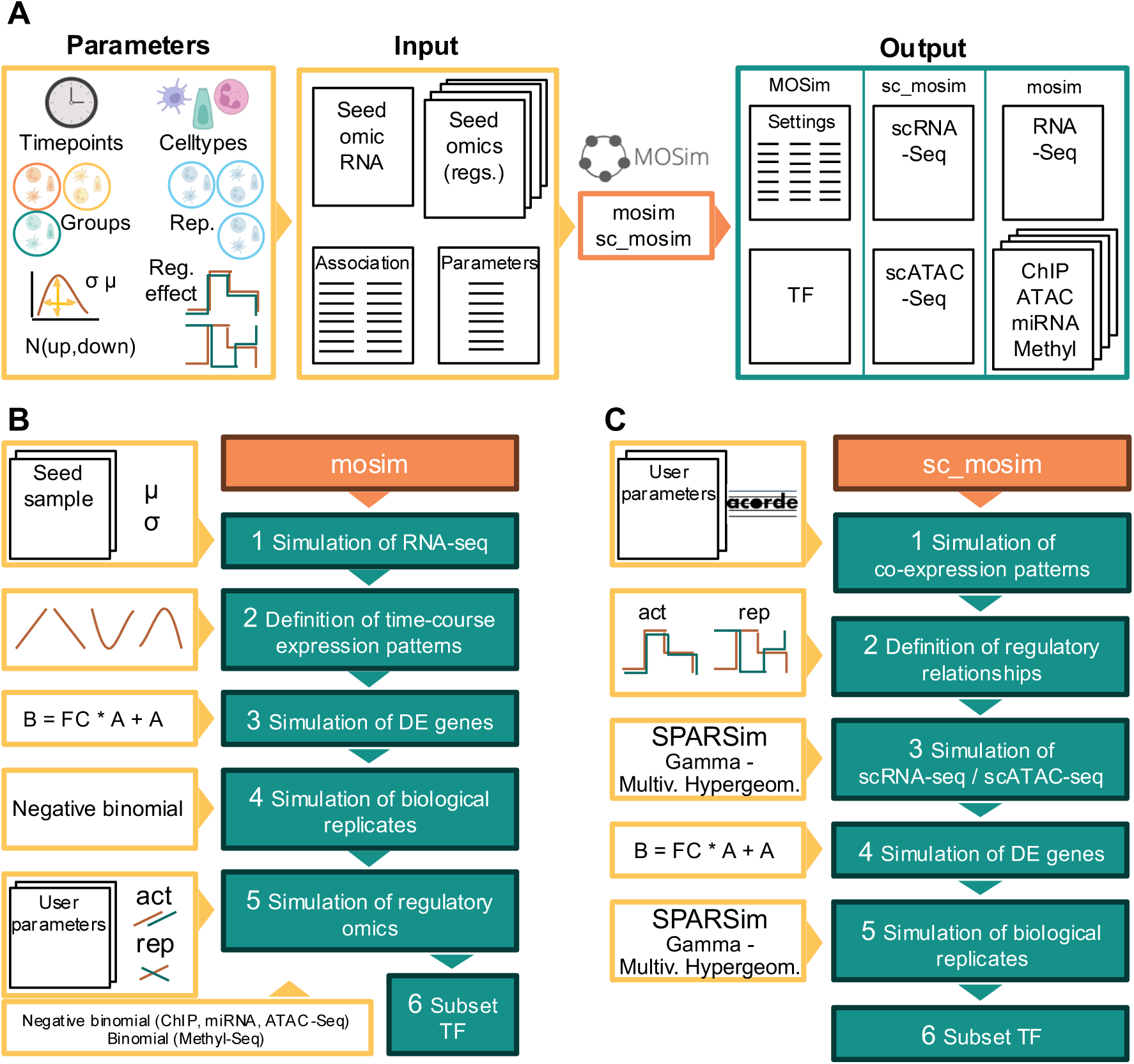
Schematic representation of the MOSim algorithms. **A)** MOSim’s simulation functionalities. **B)** Flowchart of the mosim pipeline to simulate bulk multi-omics datasets. **C)** Flowchart of the sc_mosim pipeline to simulate single-cell multi-omics datasets.

Users can define various configuration parameters related to the experimental design, such as the number of experimental groups, time-points or cell types (if applicable), replicates per experimental condition, data dispersion, number of differentially expressed genes and number of regulators with activator or repressor effects. MOSim outputs simulated count matrices for each expression and regulatory data type. Moreover, it generates a record of all parameters used in data creation (MOSim simulated settings), which is indispensable for accurately testing GRN inference bioinformatic tools (i.e. mean expression, dispersion, time profile, fold change etc.) (Figure 1A).

The MOSim package includes two main functions: mosim, for bulk datasets simulation, and sc_mosim, for single-cell datasets simulation.

The simulation results include three distinct outputs: (i) the simulated omic count data, represented as a matrix for each omic modality. Each matrix contains the same number of omic features as provided in the seed data and the number of samples, groups, cells, etc., specified by the user; (ii) for gene expression, a table listing the genes simulated as differentially expressed, along with their temporal profile (only for bulk; see Table 1); and (iii) a table for each omic modality detailing the regulatory relationships provided by the user, including the simulated activator or repression regulations (see Tables 2 and 3).

**Table 1:** MOSim-defined settings for RNA-seq simulation example. ID: Gene identifier; DE: Whether the gene is differentially expressed (TRUE) or not (FALSE); GrX: Type of gene temporal profile in experimental group X; Tmax.GrX: For transitory profiles, time point where the minimum or maximum is reached in the corresponding group X.

| ID | DE | Gr1 | Gr2 | Tmax.Gr1 | Tmax.Gr2 |
| --- | --- | --- | --- | --- | --- |
| ENSMUSG00000097082 | TRUE | Tran.Ind. | Tran.Ind. | 1.872 | 1.311 |
| ENSMUSG00000020205 | TRUE | Tran.Ind. | Cont.Ind. | 2.114 | NA |
| ENSMUSG00000055493 | TRUE | Tran.Ind. | Cont.Rep. | 3.062 | NA |
| ENSMUSG00000087802 | FALSE | Flat | Flat | NA | NA |
| ENSMUSG00000017204 | TRUE | Tran.Ind. | Cont.Rep. | 2.610 | NA |
| ENSMUSG00000017221 | TRUE | Tran.Ind. | Cont.Ind. | 1.359 | NA |
| ENSMUSG00000052726 | TRUE | Tran.Ind. | Tran.Ind. | 3.178 | 1.626 |

**Table 2:** MOSim-defined settings for ATAC-seq simulation example. ID: Genomic coordinates of ATAC-seq region (chromosome, and start and end positions for chromatin-accessible regions); Gene: Regulated gene; Effect.GrX: Regulatory effect of the ATAC-seq region on gene expression in experimental group X; GrX : temporal profile of the ATAC-seq region in experimental group X.

| ID | Gene | Effect.Gr1 | Effect.Gr2 | Gr1 | Gr2 |
| --- | --- | --- | --- | --- | --- |
| 10_111588324_111588448 | ENSMUSG00000097082 | activator | activator | Trans.Ind. | Trans.Ind. |
| 10_111588324_111588448 | ENSMUSG00000020205 | activator | NA | Trans.Ind. | Trans.Ind. |
| 10_11358301_11358431 | ENSMUSG00000055493 | activator | activator | Trans.Ind. | Cont.Rep. |
| 10_11358301_11358431 | ENSMUSG00000087802 | NA | NA | Trans.Ind. | Cont.Rep. |
| 11_98682094_98682786 | ENSMUSG00000017204 | repressor | activator | Trans.Rep. | Cont.Rep. |
| 11_98682094_98682786 | ENSMUSG00000017221 | repressor | repressor | Trans.Rep. | Cont.Rep. |
| 1_140257767_140257897 | ENSMUSG00000052726 | activator | activator | Trans.Rep. | Trans.Ind. |

**Table 3:**
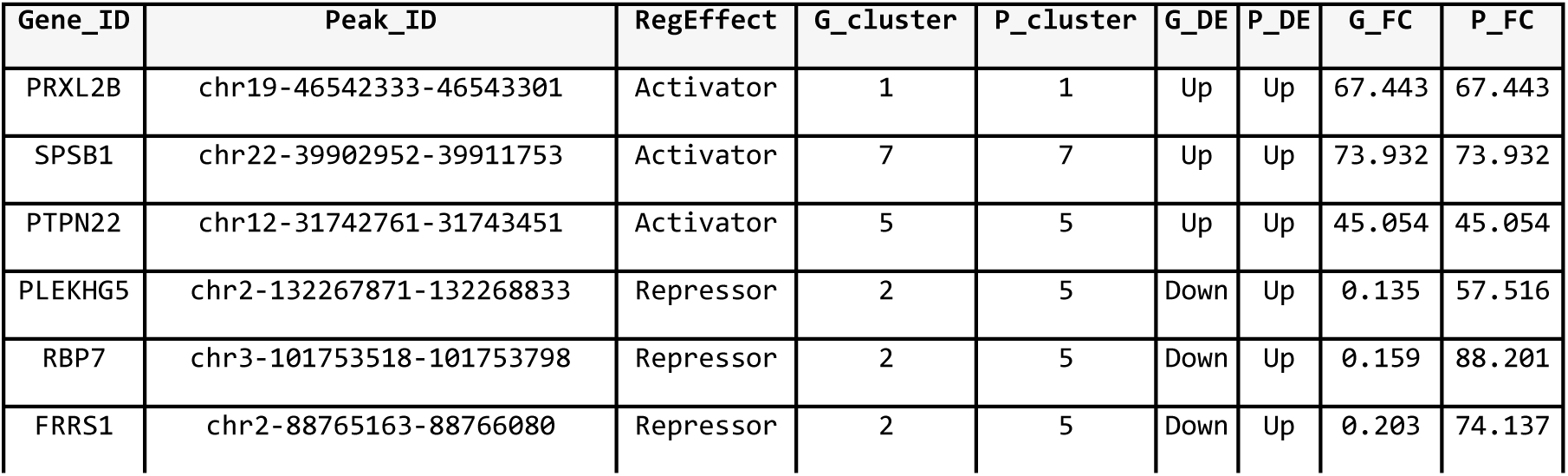
MOSim-defined settings for scRNA-seq and scATAC-seq for the simulation example. Gene_ID: Gene identifier; Peak_ID: Peak identifier; RegEffect: Regulatory effect of the scATAC-seq region on gene expression in experimental Group 2; G_cluster: gene expression profile across cell types; P_cluster: peak accessibility profile across cell types; G_DE: how the gene is differentially expressed; P_DE: how the peak is differentially accessible. G_FC: Fold Change applied to induce differential gene expression in Group 2 compared to Group 1; P_FC: Fold Change applied to induce differential peak accessibility in Group 2 compared to Group 1.

| Gene_ID | Peak_ID | RegEffect | G_cluster | P_cluster | G_DE | P_DE | G_FC | P_FC |
| --- | --- | --- | --- | --- | --- | --- | --- | --- |
| PRXL2B | chr19-46542333-46543301 | Activator | 1 | 1 | Up | Up | 67.443 | 67.443 |
| SPSB1 | chr22-39902952-39911753 | Activator | 7 | 7 | Up | Up | 73.932 | 73.932 |
| PTPN22 | chr12-31742761-31743451 | Activator | 5 | 5 | Up | Up | 45.054 | 45.054 |
| PLEKHG5 | chr2-132267871-132268833 | Repressor | 2 | 5 | Down | Up | 0.135 | 57.516 |
| RBP7 | chr3-101753518-101753798 | Repressor | 2 | 5 | Down | Up | 0.159 | 88.201 |
| FRRS1 | chr2-88765163-88766080 | Repressor | 2 | 5 | Down | Up | 0.203 | 74.137 |

In addition to the primary MOSim functions for simulating bulk (mosim) or single-cell (sc_mosim) multi-omic datasets, the package provides several other useful functions. These include functions for modifying seed data (omicData and sc_omicData), adjusting default omic parameters (omicSim), and retrieving simulation results and settings (omicResults, omicSettings, sc_omicResults and sc_omicSettings).

### Bulk multi-omic GRNs: the mosim functionality

The mosim workflow (Figure 1B) consists of the following steps:

1. As the more extended assumption for RNA-seq data, a negative binomial (NB) distribution is applied to generate a bulk RNA-seq count matrix, obtaining the mean and dispersion from the seed sample and the amount of variability across replicates set by the user.
2. Differentially expressed genes (DEGs) are randomly selected from the seed RNA-seq sample. DEGs are labelled with one of the following time-course patterns in each experimental group: continuous induction (increasing linear pattern), continuous repression (decreasing linear pattern), transitory induction (quadratic pattern with an intermediate maximum), transitory repression (quadratic pattern with an intermediate minimum), or flat, which is also the default pattern for non-DEGs.
3. Expression profiles are simulated based on the seed count values to closely reflect real data distributions. For transitory profiles, the algorithm randomly selects the time point at which the expression reaches its maximum or minimum and simulates a quadratic pattern. For continuous profiles, the algorithm randomly defines both the expression value at the first time point and the slope of change over time, simulating a linear pattern. These patterns vary depending on the coefficient values of the simulation function, particularly as the number of time points increases, and thus, although there are four theoretical temporal profiles, the simulated profiles encompass a wider range of patterns. DEGs with flat profiles or DEGs in a two-group design with no time points are modelled by introducing a fold-change in one of the experimental conditions. For designs with more than two experimental groups, the first serves as the reference and the fold-change is applied to a random selection of the remaining groups.
4. After generating gene expression values for each condition, replicates are simulated using a NB distribution.
5. All bulk MOSim data types (except Methyl-seq) are assumed to follow a NB distribution. Therefore, the NB is also used to simulate replicates for the remaining omics, but subjected to the simulated settings of the provided regulatory data and a randomly chosen direction of regulation. Regulators labelled as activators adopt the same profile as their associated genes, while repressors follow the opposite pattern.
6. For Methyl-seq, proportions are generated instead of counts based on the binomial distribution, following the strategy described in [9]. TF expression values are extracted from the simulated RNA-seq data to simulate TF regulation.

A detailed explanation of the bulk mosim algorithm implementation is provided in Supplementary File 1.

### Single-cell multi-omic GRNs: the sc_mosim functionality

The workflow of sc_mosim (Figure 1C) consists of the following steps:

1. Following the approach used by the acorde R package for defining isoform profiles across cell types in single-cell RNA-seq [10], gene expression and peak accessibility values in the seed datasets are reorganised to build synthetic features following cross-cell type patterns, i.e., indicating low or high expression in a given cell type.
2. Peak accessibility values are rearranged to reflect the regulatory relationship between scRNA-seq and scATAC-seq. Regulators labelled as activators share the same cross-cell type profile as their associated gene, while repressors have the opposite pattern.
3. Feature intensity, variability (variance of normalised counts across cells of the same cell type) and library size of the rearranged seed scRNA-seq and scATAC-seq datasets are estimated using SPARSim. A reference dataset is then simulated for each omic data type using a Gamma-Multivariate Hypergeometric model [11].
4. DEGs are randomly selected from the reference scRNA-seq. DEGs and their associated differentially accessible peaks between experimental groups are generated by introducing a fold-change in the experimental conditions, using the first condition as the reference. Additionally, random noise is added to the quantification values to introduce realistic variability between experimental groups and across features.
5. Feature intensity and library size of the simulated scRNA-seq and scATAC-seq count matrices for each experimental group are estimated using SPARSim. Biological replicates are then simulated using the Gamma-Multivariate Hypergeometric model [11], with the estimated parameters and a small random variability. TF expression values are extracted from the simulated scRNA-seq data to simulate TF regulation.

A detailed explanation of the single-cell sc_mosim algorithm implementation is provided in Supplementary File 1.

### Validation of the bulk (mosim) simulation approach

To demonstrate mosim’s capabilities for bulk sequencing data, we simulated RNA-seq and ATAC-seq data with five time points, two experimental groups, and three replicates, using the STATegra [12] samples included in the MOSim R package as seed data. We set the number of DEGs to 15% and modelled the five temporal profiles previously described. MOSim returns two types of output. The omicResults function returns a list containing the simulated data matrix for each omic, with features in rows and observations in columns. The second results object, accessible via the omicSettings function, includes the mosim-generated settings for the simulated relationships between gene expression and the rest of omics, as illustrated in Tables 1 and 2 containing simulation settings for RNA-seq and ATAC-seq, respectively. For instance, gene ENSMUSG00000052726 is identified as a DEG, displaying transitory repression in condition 1 and transitory induction in condition 2. The chromatin-accessible region 1_140257767_140257897 is simulated as a significant activator of this gene in both conditions, thereby following the same temporal profiles as the regulated gene.

We applied the K-means method to cluster simulated gene profiles, aiming to verify that the algorithm generates the expected profiles. Features with an average expression per condition of less than one count per million were filtered out. We compared the MOSim assigned profile with the average profile of the corresponding cluster and classified a gene as correctly simulated if both profiles coincided (for example, if a gene was assigned a constant induction profile and clustered with a group exhibiting a continuous increase in expression). The optimal number of clusters was found to be k = 7 for K-means clustering, which resulted in one cluster per simulated pattern and time point of maximal or minimal expression. Figure 2A displays the K-means clustering results for the simulated RNA-seq data in group 1, revealing that most genes in the cluster faithfully follow the mean cluster profile, as expected. Overall, less than 0.5% of the simulated profiles were assigned to an incorrect cluster.

**Figure 2:**
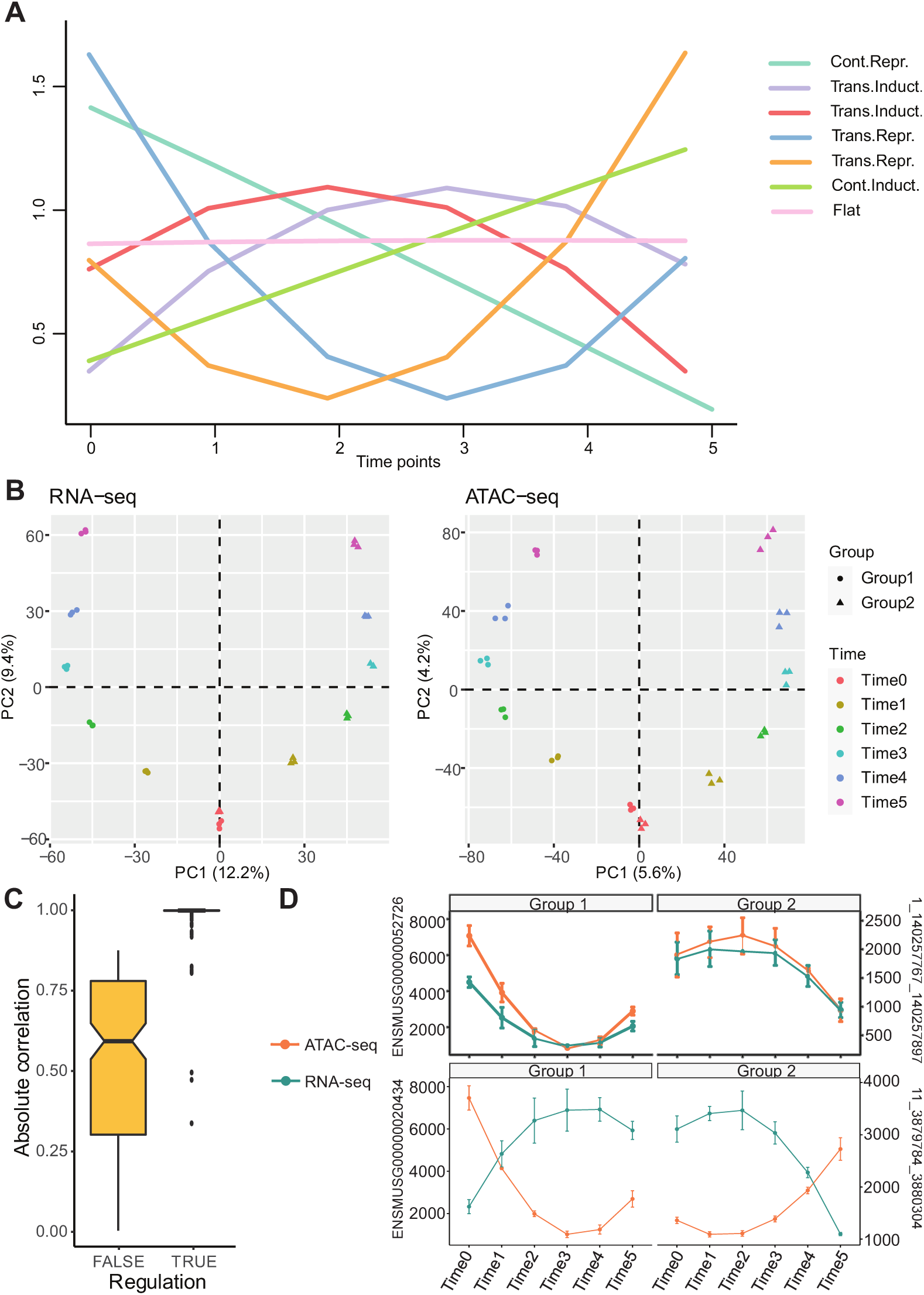
**A)** Representation of K-means clusters for bulk RNA-seq in group 1. The coloured lines represent the cluster mean profiles. The temporal simulated profiles associated with each cluster are indicated in the figure legend: one cluster corresponds to continuous repression, one to continuous induction, and one to a flat profile. Additionally, there are two clusters showing a maximum peak at different intermediate time points for transient induction, and two clusters showing a minimum peak at different time points for transient repression. **B)** Exploratory analysis using Principal Component Analysis on low-count filtered data with logarithmic transformation. The first principal component separates the samples by the experimental group, while the second summarises the temporal profile. X- and Y-axis labels indicate the percentage of variability explained by the corresponding principal component. **C)** Boxplot of absolute Pearson’s correlation values from interactions of ATAC-seq regulators with genes in Group 1. Regulation is TRUE when the regulator has been simulated to activate or repress gene expression. Regulation is FALSE for interactions where the regulator has not been simulated to affect gene expression. **D)** Two random examples of gene-regulator temporal profiles in each group. The left Y-axis shows gene expression values, while the right Y-axis shows counts for ATAC-seq regions. Vertical bars at each time point show the standard deviation of the 3 simulated replicates.

We further evaluated the simulated data using Principal Component Analysis (PCA). The PCA score plot (Figure 2B) indicates that the simulated data effectively recapitulated a quality time course dataset, where replicates were clustered together and consecutive time points were proximate.

Following the validation of individual omic data, relationships between gene expression and regulatory omics were evaluated by measuring correlations. An interaction between a regulator and a gene is expected to yield a high absolute correlation value when the regulator exerts a modelled effect on the gene, sharing the same profile type for activation or exhibiting an opposite pattern (i.e., continuous induction vs continuous repression) for repression. When no effect is modelled between the gene and regulator, the profiles will exhibit uncorrelated patterns (e.g. transitory vs continuous). Pearson’s correlations were calculated for each interaction and separately for each group. Interactions involving transitory profiles in both the regulator and the gene may include a delayed response, where the signal maxima -or minima-occur at different time points, with Pearson’s correlation failing to capture these regulatory relationships. To address these scenarios, we also computed a lagged correlation, limiting the sliding of time points to a maximum of two to control for false positives, and selecting the maximum value from Pearson and lagged correlations as the correct measure. In the ATAC-seq example (Figure 2C), 99.4% of interactions with a modelled activator or repressor effect displayed a correlation value above 0.9, while 0% of interactions without a modelled effect reached this threshold. Correlation values varied widely for these “no effect” interactions, ranging from the expected low values to relatively high ones. The latter can often be attributed to partial overlap between non-comparable profiles, such as a transient induction profile in the gene alongside a continuous induction profile in the regulator, both sharing an increasing linear trend over the same time points. This pattern aligns with the algorithm’s intended and expected behaviour. Figure 2D presents simulated temporal profiles for each experimental group, showcasing two randomly selected gene-regulator pairs. In the first pair (top plots), the regulation is activation in both groups, while in the second pair (bottom plots), the regulation is repression, also consistent across both groups.

### Validation of the single-cell (sc_mosim) simulation approach

To demonstrate the utilities of sc_mosim for single-cell sequencing data, we simulated scRNA-seq and scATAC-seq data with six cell types, two experimental groups, and three replicates. We used the pbmcMultiome dataset available from SeuratData [13] as seed data and the generegulator association list provided in the MOSim R package. We set the number of DEGs to 30% upregulated and 20% downregulated. Variances were set to 0.1 between replicates and 0.3 between experimental groups, and we allowed for the modelling of co-expression patterns across cell types, following seven random profiles. Finally, we defined 20% activator and 10% repressor regulators in Group 1, and 10% activators and 20% repressors in Group 2.

In single-cell simulations, MOSim generates two main types of output. The sc_omicResults function retrieves a list containing the simulated data matrices for each omic, experimental group and biological replicate, with features in rows and cells in columns. The second results object, extracted with the sc_omicSettings function, includes the MOSim-generated settings that associate genes and peaks (Table 3), and specify TFs with their target genes, along with the type of regulatory relationship between them. For example, gene PTPN22 is identified as an upregulated DEG that follows the across-cell-type expression pattern 5 (Figure 3A). The chromatin-accessible region chr12-31742761-31743451 is modelled as a significant activator of this gene, following the same across-cell-type profile as the regulated gene. Conversely, the association between the gene RBP7 and chromatin-accessible region chr3-101753518-101753798 exemplifies a repressor effect of the regulator omic, where gene and peak follow opposite patterns (clusters 2 and 5, respectively), with the gene downregulated when the regulator is upregulated (Table 3).

**Figure 3:**
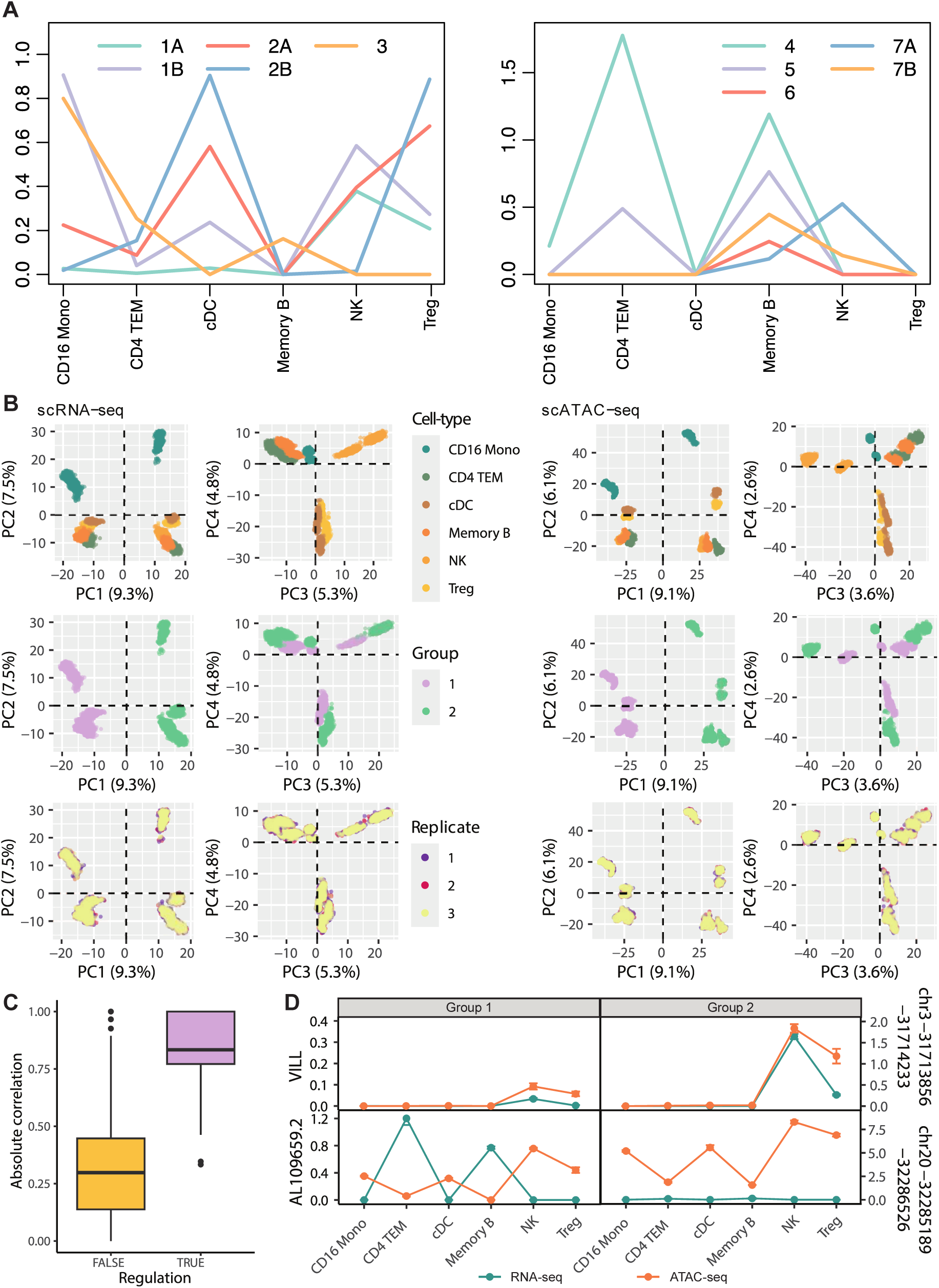
**A)** Representation of clustering patterns for single-cell RNA-seq across cell types for group 1; split into two plots to improve visualization and cluster differentiation. The coloured lines represent the cluster mean profiles. **B)** Exploratory analysis using Principal Component Analysis to visualize clustering of cells coloured by cell type, experimental group and replicate. X- and Y-axis labels indicate the percentage of variability explained by the corresponding principal component. **C)** Boxplot of absolute Kendall correlation values from interactions of scRNA-seq genes in Group 1 with scATAC-seq regulators. Regulation is TRUE when the regulator has been simulated to activate or repress gene expression. Regulation is FALSE for interactions where the regulator has not been simulated to affect gene expression. **D)** Two examples of gene-regulator single-cell simulated profiles in each group. The left Y-axis shows gene expression values, while the right Y-axis shows counts for scATAC-seq regions. Vertical bars at each time point show the standard error of the mean of the cells for the 3 simulated replicates.

To demonstrate the robustness of the single-cell MOSim framework for GRN simulation, we assessed its capacity to generate the expected across-cell-type expression profiles. Single-cell data is typically characterised by a high abundance of zeros and many cells belonging to the same cell type, leading to increased noise and outliers. Given the robustness of Spearman’s correlation distance and K-medoids clustering techniques in noisy scenarios, we used them to extract and cluster the simulated feature profiles across cell types. The cluster average profiles were then compared to the sc_mosim simulated profiles after excluding genes with flat expression profiles. A feature was deemed correctly simulated if both profiles matched. To achieve this, we set the optimal number of clusters to K = 10 for K-medoids clustering, which resulted in one or two clusters per simulated co-expression pattern, minus flat expression. Clustering of the simulated scRNA-seq and scATAC-seq revealed that most features closely adhered to the mean cluster profiles as expected (Figure 3A), with only 3.3% of simulated profiles assigned to an incorrect cluster.

We further assessed whether cells from the same cell types, experimental groups, and biological replicates clustered according to the defined simulation settings using PCA for dimensionality reduction (Figure 3B). PCA results showed robust clustering of the simulated data, capturing a high-quality single-cell dataset where PC1 separated cells by experimental group, while PCs 1 to 4 represented the cohesive clustering of cell types (Figure 3B). Additionally, while the majority of data variability was due to differences specified between groups, small variability between biological replicates was also observable (Figure 3B).

To evaluate whether simulated regulatory relationships presented stronger correlations than non-regulatory peak-gene associations, Kendall’s correlations between gene and peak profiles were computed within each simulated experimental group. A strong absolute correlation is expected for pairs when a regulatory effect was modelled, reflecting similar activation or opposite repression profiles. In contrast, non-regulatory peak-gene interactions typically display lower and more variable correlation values due to differences in absolute terms. As shown in Figure 3C, 79.5% of interactions with modelled activator or repressor effects had absolute correlation values exceeding 0.7, while “no effect” interactions displayed a broader range, centered at 0.32 absolute correlation. This range is likely due to partial overlaps, such as shared trends between cell types, which are expected outcomes of the simulation.

Finally, Figure 3D illustrates simulated feature profiles across cell types for two pairs of gene-regulator associations, one with an activator effect and the other with a repressor effect. The first regulation (top plots) represents activation in both groups, whereas the second regulation (bottom plots) represents repression across both groups.

### Simulation of multilayered Gene Regulatory Networks

Finally, we illustrate how MOSim effectively simulates multilayered GRNs. Simulating GRNs is challenging due to the complex many-to-many relationships among some regulators and their target genes. For example, a TF or microRNA might regulate multiple target genes with varying regulatory relationships, while the same gene could be influenced by multiple factors. A multimodal GRN simulation algorithm must therefore produce a consistent dataset with expression patterns reflecting these different regulatory patterns. In MOSim, users can specify a desired percentage of active regulatory relationships, and the algorithm adjusts regulatory pairs and profiles to achieve this level of regulation across layers (Figures 2 and 3).

To demonstrate MOSim’s capabilities in modelling multilayered regulatory interactions, we used the STATegra dataset [12] to simulate RNA-seq, miRNA-seq, and TF data. The simulation was performed with a sequencing depth of 30 million reads, two experimental groups, three replicates per group, and six time points, forming a detailed experimental design. Additionally, we specified that 5% of genes be differentially expressed, and 40% of miRNA-seq over the total number of regulators should be repressor effects.

Given the complexity of visualising the simulated GRN, we selected the first 100 differentially expressed genes and plotted their corresponding GRNs for each experimental group (Figures 4A and 4B). To illustrate the profiles of features in these simulated subnetworks and the efficiency of MOSim in creating consistent expression patterns across different layers, we generated heatmaps for each experimental group (Figure 4C). To facilitate visualisation and interpretation, we calculated the mean expression across replicates for each time point and experimental group, scaling the expression values across modalities, since each omic layer may have different value ranges. Figure 4C demonstrates MOSim’s capacity to simulate distinct feature profiles across layers, accurately reflecting both activator and repressor regulatory effects.

**Figure 4:**
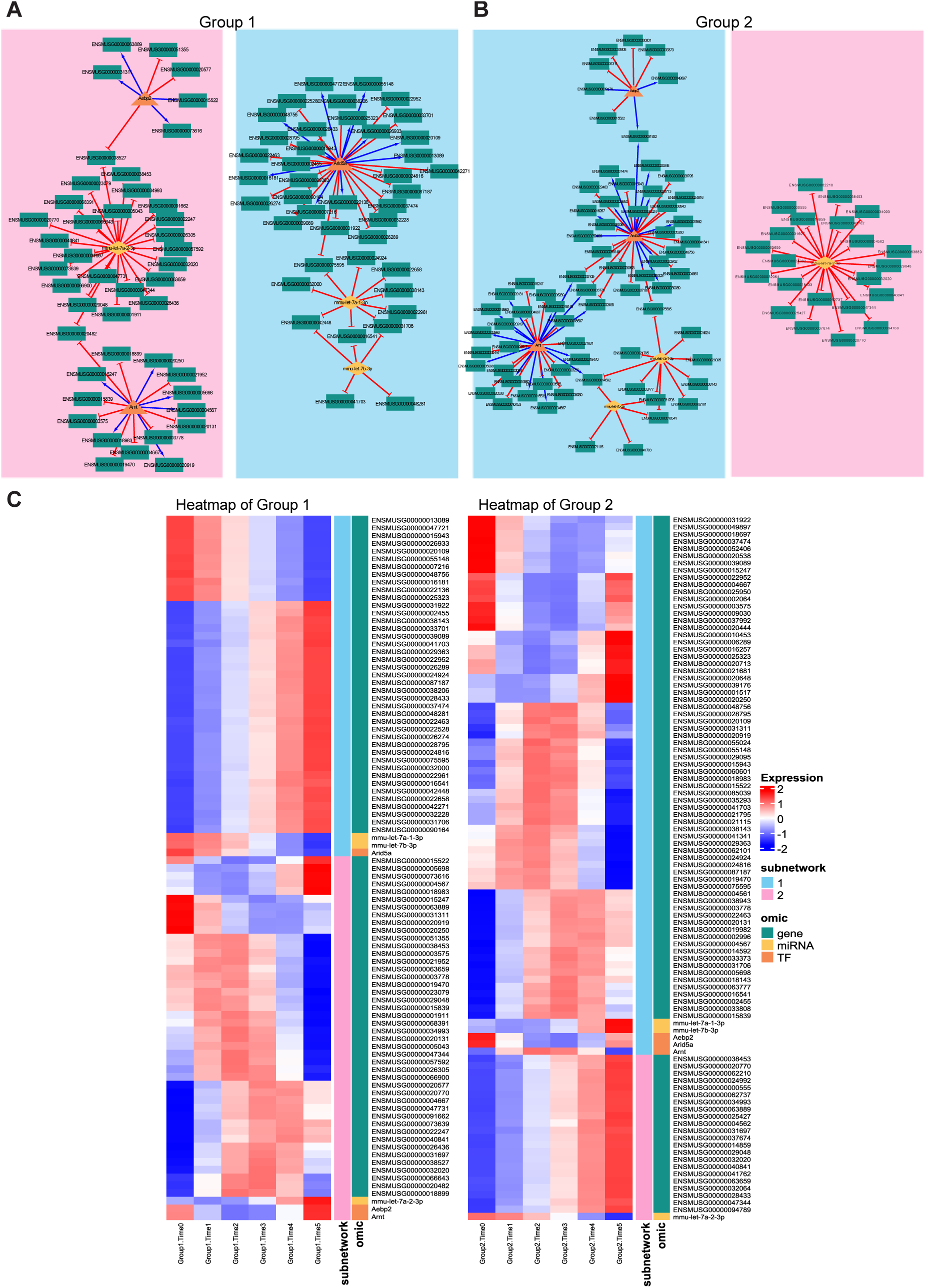
Representation of multi-layer regulatory networks simulated by MOSim. Genes are represented in green, transcription factors are in orange, and miRNAs are in yellow. Blue arrows represent activator regulations, while red arrows repressor regulations. **A)** Gene Regulatory Network for Group 1. **B)** Gene Regulatory Network for Group 2. **C)** Heatmaps for the expression profiles of the genes, miRNAs and Transcription Factors in Gene Regulatory Networks of Groups 1 and 2. The right Y-axis shows the omic data type and the subnetwork they belong to (which refers to the connected subnetworks observed in A) and B) framed in pink and blue rectangles).

This example demonstrates that MOSim can generate consistent, complex modules with both positive and negative regulatory relationships, spanning multiple layers and including one-to-many and many-to-many interactions—providing a unique capability to simulate the complexity of gene regulation.

### Application of MOSim for benchmarking a GRN inference tool

To demonstrate one potential application of MOSim simulations, we used MOSim-generated data to test MORE (Multi-Omics Regulation), a tool designed to infer GRNs from bulk multi-omics data [14]. Specifically, we simulated RNA-seq, miRNA-seq and TF data with MOSim using the STATegra dataset [12]. The simulation was configured with a sequencing depth of 30 million reads, two experimental groups, 20 time points per group, and one replicate per time point. Additionally, we set the percentage of differentially expressed genes to 50%, and the percentage of significant regulations to 60%.

Prior to applying MORE, the RNA-seq count matrix was pre-processed. Low-count genes were filtered out with the NOISeq R package [15], using a threshold of 1 count per million. Count data was normalized with the weighted trimmed mean of M-values (TMM) normalisation in the NOISeq package and voom-transformed [16]. Differential expression analysis between groups 1 and 2 was performed with the limma R package [17], yielding 10593 DEGs (FDR < 0.05). These DEGs were set as the target omic features required by MORE. The miRNA-seq and TF data were used as the regulatory omics. For GRN inference, we applied the MORE PLS1 option with auto-scaling and Jack-Knife resampling for the selection of significant regulators.

MORE fitted 5573 models, one for each gene with potential regulators. The MOSim simulation provided a total of 370,566 potential regulatory interactions (gene-regulator pairs), 47% of which were simulated as significant in at least one of the groups (174,051 in group 1 and 174,067 in group 2). These significant regulations served as the ground truth, or positive instances, to evaluate MORE’s performance. At a significance level of 0.05, MORE identified 233,598 significant regulations in group 1 and 240,474 in group 2 that were compared to the positive instances. The analysis yielded similar error metrics for both groups, with a slightly better performance observed in miRNA-seq compared to TFs. Overall, MORE achieved a sensitivity of 85.5% and an F1-score of 62.9%. These results demonstrate MORE’s ability to detect significant regulatory interactions, while also indicating areas where the tool could be improved or where hyperparameter tuning might enhance its performance.

This example highlights how MOSim can serve as a reliable ground truth framework for evaluating the performance of GRN inference tools during their development.

### Benchmarking scMOSim’s scRNA-Seq simulations using a deep learning algorithm

To further demonstrate other applications of MOSim simulations, we tested it using a Variational Autoencoder (VAE)-based tool. VAEs are capable of learning meaningful latent representations of single-cell data. Unlike standard autoencoders, VAEs impose a probabilistic structure on the latent space, enabling more robust feature extraction and better generalization across datasets. This makes VAEs particularly useful for clustering, dimensionality reduction, and transcription factor perturbation analysis[18]. Examples of VAE models for single-cell data include scGen[19], VEGA[20], siVAE[21], scVAE[22], scDHA[23], scVI[24], manatee[25] and ScInfoVAE[26].

We tested scMOSim-generated single-cell RNA-Seq data using the VAE-based tool, single-cell Decomposition using Hierarchical Autoencoder (scDHA)[23]. scDHA first removes noise using a non-negative kernel autoencoder and then projects the data into a low-dimensional space using a stacked Bayesian autoencoder. Finally, it applies iterative perturbations to reduce overfitting and create a more generalized representation.

We used one replicate from a single experimental group of scRNA-Seq data simulated with scMOSim to evaluate cell clustering with scDHA. The clustering identified five of six simulated cell types, with one cluster combining cDC and Treg cells (Table 4). The Adjusted Rand Index (ARI) score was 0.949, showing high agreement between predicted and true labels.

**Table 4:**
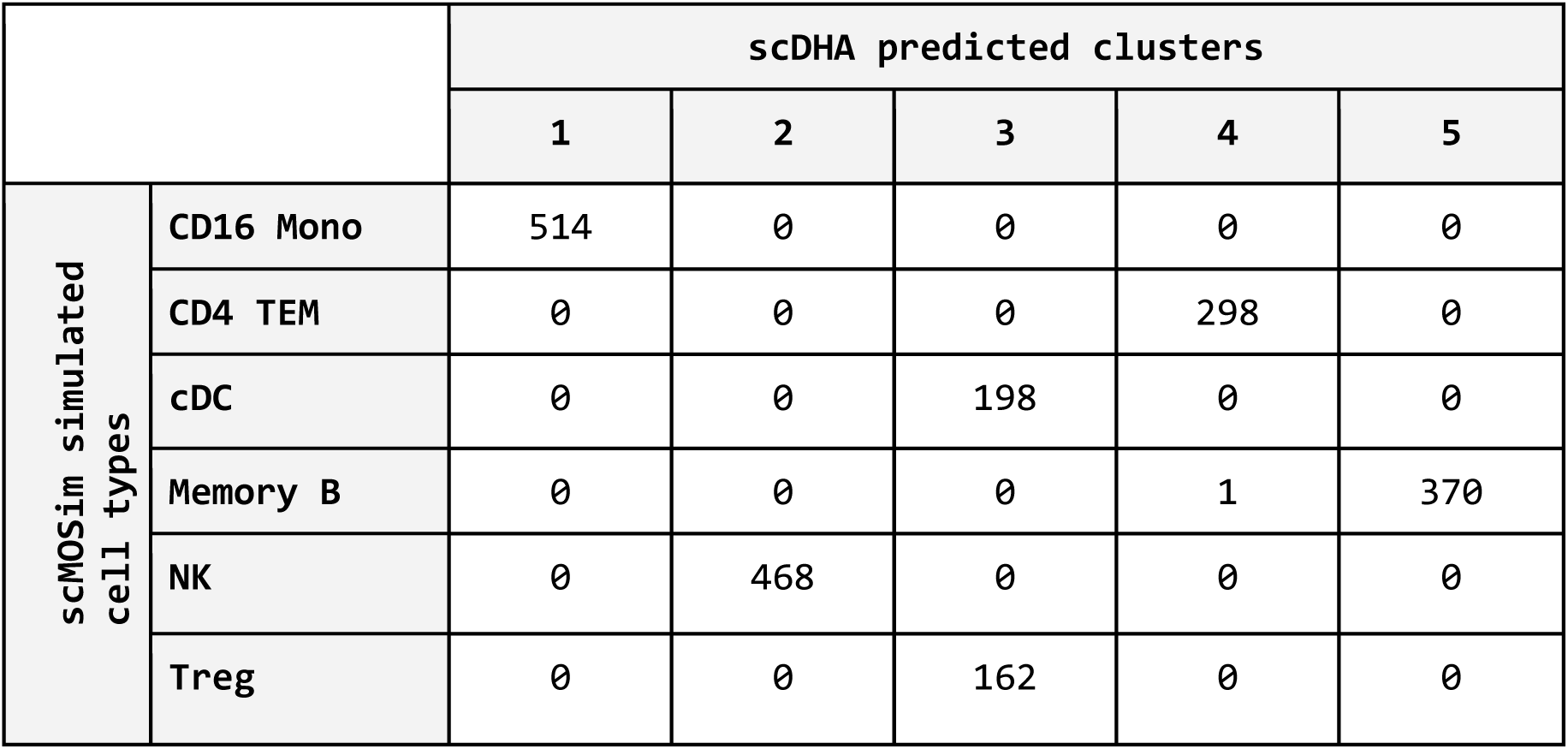
Number of cells per cell-cluster identified using scDHA, compared with ground truth cell type groups simulated using scMOSim.

These results demonstrate scMOSim’s, and its underlying algorithm SPARSim’s[11], ability to reliably simulate single-cell RNA-Seq ground truth datasets with different cell-types sufficiently distinguishable as to be identified by a VAE algorithm such as scDHA.

## Discussion

Multi-omic assays, facilitated by massively parallel sequencing technologies, have greatly enhanced our ability to profile regulatory mechanisms in biological systems [1,2], leading to a deeper understanding of diseases and model organisms. However, benchmarking studies of bioinformatic tools designed to elucidate multi-layered GRNs by integrating multi-omics datasets have exposed notable discrepancies in library preparation strategies and analysis methods [27]. These discrepancies underscore the complex challenge of accurately identifying GRNs. As multi-omic sequencing continues to gain traction in the study of regulatory mechanisms, there is a pressing need for tools that support rigorous GRN inference assessment.

MOSim was developed to provide a robust framework for simulating bulk and single-cell multi-omics data in a controlled setting. Using a seed dataset and a regulator-gene association matrix, MOSim generates realistic simulated count matrices for both bulk and single-cell transcriptomics data, as well as for associated regulatory omics. For bulk data, the simulation is based on the negative binomial distribution, while for single-cell data, it leverages the well-established simulator SPARSim[11]. By using a seed dataset as a reference to infer distributions, MOSim generates count matrices that closely mirror real omic data, offering a more authentic representation than simulators that artificially construct count matrices without a real-data foundation [28].

Additionally, MOSim operates at the count matrix level rather than simulating read data, providing a unified framework for generating multi-omics data across different library preparation methods (e.g., SmartSeq2, 10x Genomics). This allows users to select a preferred method as the seed dataset for MOSim, adding flexibility to the simulation process.

MOSim enables a fast and effortless generation of bulk and single-cell count data matrices for multiple omic types, supporting flexible experimental designs. Importantly, the algorithm can simulate complex regulatory relationships between gene expression and other molecular components, guided by prior knowledge, such as target mRNA-microRNA associations. This flexibility in defining experimental designs, DEGs, and active regulators makes MOSim a versatile tool for a variety of different applications, including: i) validating methods aimed at modelling complex, multi-layered regulatory networks, ii) benchmarking multi-omics data integration pipelines, iii) benchmarking GRN inference tools [2], iv) evaluating differential expression and accessibility analysis tools [24], v) testing single-cell data clustering methods (Supplementary File 1) [24], vi) evaluating multi-omics visualization tools, vii) testing methods for time-series analysis in RNA-seq data [29], among others. Several tools have already been tested using MOSim simulations, including DEGRE [30], scAI [31], JISAE [32], GR-NIC [33] and scLRTD [34], highlighting MOSim’s ability to specify an association matrix for linking regulators with transcripts further allows users to tailor MOSim outputs to align with the intended integration goals of their analysis tools.

The MOSim framework has some limitations. Currently, single-cell simulation is restricted to scRNA-seq and scATAC-seq, as these are presently the only two commercially available sequencing techniques that can be simultaneously performed on the same cell. As additional single-cell omics techniques become widely available, extending MOSim to other data types will be straightforward based on its bulk framework. At this point, MOSim is not prepared to simulate GRN with spatial resolution, which could be inferred from spatial multi-omics data. While these datasets are not yet widespread, they might be in the near future. We envision that the flexible MOSim simulation framework could incorporate the spatial information either as covariates of the regulatory model or by modelling cell-to-cell communication signals as an additional regulatory layer. These possibilities are to be explored in future work. Finally, both bulk and single-cell modules are designed to simulate gene regulatory relationships based on sequencing data, limiting applicability to other omics layers like proteomics and metabolomics, which may influence gene regulation in more complex or uncertain ways. Future work will also explore extending MOSim to simulate interactions between gene expression, the proteome, and the metabolome.

## Conclusion

The integration of multi-omics datasets for GRN identification remains a challenging task. We demonstrate that MOSim serves as an essential resource for benchmarking integration tools, filling a critical gap in the multi-omics sequencing field.

## Methods

The MOSim algorithms are introduced in the results section and extended in Supplementary File 1. The algorithms are implemented in R and mainly use R packages dplyr [35], purrr [35], Stats [36], Iranges [37], Seurat [38], SPARSim [11], and adapted scripts from Acorde [10] and WGBSSuite [9].

### MOSim algorithms assessment

The performance of the MOSim bulk simulation was tested with mouse multi-omics data from the STATegra project [12], while single-cell simulation performance was evaluated using the human pbmc.multiome 10x Genomics dataset from the SeuratData R package [13].

For the bulk data, K-means clustering [39] was applied to the simulated feature profiles to assess the correct simulation of temporal expression patterns. For single-cell data, the simulated count matrix was aggregated to obtain the average count per cell type. Spearman’ distance (1 - Spearman’s correlation [40]) and partition around medoids (K-medoids [41]) clustering were then used to cluster gene expression profiles across cell-types. In both cases, the optimal number of clusters was obtained by combining the maximisation of Silhouette’s coefficient and minimising the intra-cluster variability.

In both bulk and single-cell simulations, a log transformation (*log*(*x* + *1*))[42] was applied to the data. PCA was used to confirm that clustering aligned with the simulation settings. Finally, to validate gene-regulator relationships, Pearson’s correlation was computed for bulk data and Kendall’s T_b_ correlation for single-cell data [43]. These correlations were compared with 20.000 random feature pairs with no simulated regulatory effects.

## Availability of data and materials

The package is released under the GNU Public License to the community as a package named MOSim, for Multi-Omics Simulator, at Bioconductor (https://bioconductor.org/packages/MOSim/).

Bulk example data in MOSim was generated by the STATegra project [12]. Single-cell example data is available in the pbmc.multiome dataset in the SeuratData R package [13]. Code to reproduce the figures in the manuscript is available on github (https://github.com/BiostatOmics/MOSim_plots).

## Acknowledgements

The authors thank Arianna Febbo for preliminary analyses. The development of the tool and test simulations were performed on the high-performance computing cluster Garnatxa at the Institute for Integrative Systems Biology (I2SysBio), I2SysBio is a mixed research centre formed by the University of Valencia (UV) and Spanish National Research Council (CSIC).

## Author contributions

S.T., A.C. and C.M.M. conceptualised and designed the mosim approach. S.T., A.C., C.M. and A.A.L. conceptualised the sc_mosim approach. C.M.M. developed and implemented mosim. C.M. developed and implemented sc_mosim and contributed to implementing mosim. C.M.M, C.M., M.A. and S.T. performed the analysis and generated visualisations. A.A.L. contributed to implementing sc_mosim. S.T. and A.C. envisioned the study and supervised the work. C.M., C.M.M., A.C. and S.T. drafted the manuscript. All authors read and approved the final manuscript.

## Author Biographies

Carolina Monzó is a postdoctoral researcher at the Institute for Integrative Systems Biology, Spanish National Research Council (CSIC-UV). Her research interests include bioinformatics, transcriptomics and biology of aging.

Maider Aguerralde is a PhD student at the Universitat Politècnica de València. Her research interests include method development, bioinformatics, statistics and multi-omics.

Carlos Martínez-Mira is a senior bioinformatics software engineer in Biobam Bioinformatics. His research interests include bioinformatics, statistics and software development.

Ángeles Arzalluz-Luque was a PhD student at the Institute for Integrative Systems Biology and Universitat Politècnica de València. Currently, she is a postdoctoral fellow at the Institute of Neuroscience, Spanish National Research Council (CSIC-UMH). Her research interests include bioinformatics, single cell and long-reads transcriptomics analysis.

Ana Conesa is a research professor at the Institute for Integrative Systems Biology, Spanish National Research Council (CSIC-UV). Her research interests include method development, bioinformatics and transcriptomics.

Sonia Tarazona is an associate professor at the Universitat Politècnica de València. Her research interests include method development, bioinformatics, statistics and multi-omics.

