## Supplementary File for "MOSim: bulk and single-cell multi-layer regulatory network simulator"

### Supplementary File 1

#### Bulk MOSim algorithm implementation

Bulk data simulation uses mosim, a wrapper function that performs tasks in seven stages:

(i) S*imulated-settings* stage: Genes are randomly classified as differentially expressed (labelled as "flat", "continuous induction", "continuous repression", "transitory induction" or "transitory repression") or as non-differentially expressed (labelled as "flat") (Figure 2A).

(ii) G*roups* stage: The seed RNA-seq dataset is duplicated as many times as the number of experimental groups, with each copy modified by multiplying a vector of fold change values to represent expression changes relative to the reference (Group 1). To maintain uniform sequencing depth across samples, any count changes generated for differential expression are added or subtracted from the overall sample depth.

(iii) T*ime series* stage: For each gene (g) classified as differentially expressed, time points are generated as follows:

Transform the vector for time $t\,(t = t_1, t_2, t_3, t_4)$ into $t^* = 0{,}1{,}2{,}3$.

Let $f(t^*)$ be a function of time $(t^*)$ that represents the different temporal profiles:

- $f(t^*) = a_1 + b_1 t$, for continuous induction,
- $f(t^*) = -a_1 - b_1 t$, for continuous repression,
- $f(t^*) = a_2 + b_2 t + c_2 t_2$, for transitory induction,
- $f(t^*) = -a_2 - b_2 t - c_2 t_2$, for transitory repression.

Coefficients $a_1, b_1, a_2, b_2$ and $c_2$ are computed to fall within [0,1] for induction profiles and [-1,0] for repression profiles. Thus, $a_1 = 0$ and $b_1 = \frac{1}{max\{t*\}}$. For transitory profiles, the time point $t_{max}$ (or $t_{min}$) marking the maximum (or minimum) is randomly selected within the interval $[\frac{1}{max\{t*\}} * 0.25, \frac{1}{max\{t*\}} * 0.75]$, ensuring the peak occurs within the central 50% of the timeline. The transitory coefficients are set as follows:

- If $t_{max} \geq \frac{1}{max\{t*\}}/2$: $a_2 = 0$; $b_2 = \frac{2}{t_{max}}$; and $c_2 = \frac{-1}{t_{max}^2}$
- If $t_{max} < \frac{1}{max\{t*\}}/2$: $a_2 = 1 + c_2 t_{max}^2 = 1 - t_{max}^2 / \left(\frac{1}{max\{t*\}} - t_{max}\right)^2$ ; $b_2 = -2c_2 t_{max} = 2t_{max} / \left(\frac{1}{max\{t*\}} - t_{max}\right)^2$; and $c_2 = -1 / \left(\frac{1}{max\{t*\}} - t_{max}\right)^2$

(iv) R*eplicates* stage: Gene expression for each replicate is generated using a negative binomial distribution from the stats R package [1]. For each gene (*g*) in group (*G*) at time (*t*), the mean of the distribution is $\mu_g = \max\{0.1, x_g^{t,G}\}$), and variance $\sigma_g^2 = \max\{0.03; 10^a * (\mu_g + 1)^b - 1\}$.

(v) R*egulator simulation* stage: Regulators are assigned one of three roles: activator, repressor, or "no effect". Activator count values follow the same profiles as their target genes, while repressors follow the opposite profiles (e.g. a gene with a continuous induction profile is activated by a regulator with a similar induction profile or repressed by one with a continuous repression profile). Stages *time series* and *replicates* are then rerun to simulate temporal profiles and replicates for regulators.

(vi) T*ranscription factors* stage: TF expression values are extracted from the simulated RNA-seq count table. Each TF's regulatory role (activator, repressor, or "no effect") is determined based on the temporal profile assigned to it and its associated target genes.

(vii) M*ethylation* stage (optional): If methylation data simulation is requested, methylation is not simulated from a seed dataset but instead uses adapted scripts from WGBSSuite tool [2]. CpG sites are grouped into blocks by proximity, sharing similar methylation profiles and regulatory effects. Methylation values are generated for each experimental group, adjusted by block and constrained to the 0 to 1 range. Replicates are simulated using a binomial distribution, with results presented as β values (proportions of methylation) or M values (log-transformed proportions), rather than count values as with other omics.

### Single-cell MOSim algorithm implementation

Single-cell data simulation in MOSim is executed through the sc_mosim wrapper function, which organises the process into six stages:

(i) S*imulated-settings* stage: A settings data frame is created to specify which genes and peaks will be up- or downregulated, as well as to define which genes will be activated or repressed by specific peaks.

(ii) C*lustering* stage: Gene expression patterns are simulated according to the number of user-defined profiles, with each profile represented as a logical vector (where true indicates high expression and false indicates low expression). To fit these high/low expression patterns across cell types, values are rearranged using the Acorde tool [3]. For peaks with activator or repressor functions, accessibility values are adjusted to align maximally with the regulated gene's expression. This is achieved by computing the slope of change between consecutive cell types for all genes and peaks, assigning the peak accessibility values with the smallest slope difference to the corresponding regulator of the gene.

(iii) E*stimation of parameters* stage: Using SPARSim's estimate_paramater_from_data wrapper function [4], key parameters are estimated as follows:

- Gene expression: The average of normalised counts across cells of the same cell type.
- Expression variability: Dispersion of normalised counts across cells using edgeR dispersion [5]
- Library size: The total count for each cell.

(iv) S*imulation* stage: using the estimated parameters, the single-cell simulator SPARSim is called first to simulate gene and peak quantifications with the following distributions:

- Gene or peak count distribution: Defined using a multivariate hypergeometric distribution $Y_{gc} \sim Mult.Hyper.(n = L_c, m = X_c)$, where $Y_{gc}$ is the count for gene (or peak) $g$ in cell $c$, $L$ is the estimated library size, and *X* the gene expression level.

- Biological variability distribution: Modelled by a gamma distribution [4] $X_{gc} \sim Gamma(shape = \frac{1}{\Phi_g}, scale = Z_g \Phi_g)$, where $X_{gc}$ is the count for gene (or peak) $g$ in cell $c$, $\Phi$ represents the dispersion parameter estimating cell-to-cell variability, and $Z$ is the average expression of the gene (or peak). SPARSim has shown robust ability to simulate realistic single-cell count data [4], and was ranked among the most accurate and versatile simulators in a benchmarking study [6].

(v) R*eplicates and groups* stage: Changes in count data between conditions are simulated by adjusting a fold change value for each gene within the Gamma-hypergeometric model. Additionally, biological variability between samples is included by modifying the variability parameter for each sample's model. The fold-change value per gene and the biological variability are defined through random values generated from a uniform distribution within established boundaries—either the max/min fold change values or zero and the simulated variability value from the settings stage.

(vi) T*ranscription factors* stage: Finally, TF expression values are extracted from the simulated scRNA-seq count table. Each transcription factor's regulatory role (activator, repressor, or "no effect") is determined based on its association with genes in each simulated cluster.

**Benchmarking scMOSim's scRNA-Seq simulations using a deep learning algorithm**

Single-cell omics datasets are characterized by being high-dimensional, sparse, and noisy, presenting challenges for traditional statistical and machine learning approaches. Deep learning is a powerful tool for analyzing such data, as it can effectively model complex, non-linear relationships while reducing dimensionality and denoising data [7]. In particular, variational autoencoders (VAEs) are capable of learning meaningful latent representations of single-cell data. Unlike standard autoencoders, VAEs impose a probabilistic structure on the latent space, enabling more robust feature extraction and better generalization across datasets. This makes VAEs particularly useful for clustering, dimensionality reduction, and transcription factor perturbation analysis [7]. Some examples of VAE models designed for single-cell data include scGen [8], VEGA [9], siVAE [10], scVAE [11], scDHA [12], scVI [13], manatee [14] and ScInfoVAE [15].

To demonstrate the applicability of  scMOSim-generated single-cell RNA-Seq data, we tested it using the VAE-based tool single-cell Decomposition using Hierarchical Autoencoder (scDHA) [12]. scDHA first uses a non-negative kernel autoencoder to remove noise, defined as genes or components with insignificant contributions to the part-based representation of the data. Second, it uses a stacked Bayesian autoencoder to project the data onto a low-dimensional space. Finally, it applies iterative perturbations to the low-dimensional state to mitigate overfitting and learn a generalized representation of the data.

We used one replicate from a single experimental group of scRNA-Seq data simulated with scMOSim, as described in the results section "Validation of the single-cell (sc_mosim) simulation approach", to evaluate cell clustering using scDHA. We applied the scDHA approach to cluster the simulated cells and compared the resulting clusters to the ground truth cell types defined by the scMOSim simulation. The clustering performance was high: among the six simulated cell types, scDHA identified five clusters - four of which corresponded to distinct cell types, while one cluster included both cDC and Treg cells (Supplementary Table 1). To quantitatively assess clustering quality, we computed the Adjusted Rand Index (ARI), obtaining a score of 0.949, indicating high agreement between predicted and true labels.

These results demonstrate scMOSim's, and its underlying algorithm SPARSim's [4], ability to reliably simulate single-cell RNA-Seq ground truth datasets with different cell-types sufficiently distinguishable as to be identified by a VAE algorithm such as scDHA.

**Supplementary Table 1:** Number of cells per cell-cluster identified using scDHA, compared with ground truth cell type groups simulated using scMOSim.

| | | scDHA predicted clusters | | | | |
|---|---|---|---|---|---|---|
| | | **1** | **2** | **3** | **4** | **5** |
| **scMOSim simulated cell types** | **CD16 Mono** | 514 | 0 | 0 | 0 | 0 |
| | **CD4 TEM** | 0 | 0 | 0 | 298 | 0 |
| | **cDC** | 0 | 0 | 198 | 0 | 0 |
| | **Memory B** | 0 | 0 | 0 | 1 | 370 |
| | **NK** | 0 | 468 | 0 | 0 | 0 |
| | **Treg** | 0 | 0 | 162 | 0 | 0 |


1. . R: A Language and Environment for Statistical Computing : Reference Index. 2010;
2. Rackham OJL, Dellaportas P, Petretto E, et al. WGBSSuite: simulating whole-genome bisulphite sequencing data and benchmarking differential DNA methylation analysis tools. Bioinformatics 2015; 31:2371–2373
3. Arzalluz-Luque A, Salguero P, Tarazona S, et al. acorde unravels functionally interpretable networks of isoform co-usage from single cell data. Nat. Commun. 2022; 13:1828
4. Baruzzo G, Patuzzi I, Di Camillo B. SPARSim single cell: a count data simulator for scRNA-seq data. Bioinformatics 2020; 36:1468–1475
5. Robinson MD, McCarthy DJ, Smyth GK. edgeR: a Bioconductor package for differential expression analysis of digital gene expression data. Bioinformatics 2010; 26:139–140
6. Cao Y, Yang P, Yang JYH. A benchmark study of simulation methods for single-cell RNA sequencing data. Nat. Commun. 2021; 12:6911
7. Brendel M, Su C, Bai Z, et al. Application of Deep Learning on Single-cell RNA Sequencing Data Analysis: A Review. Genomics Proteomics Bioinformatics 2022; 20:814–
